# Levodopa reduces consumption of multiple classes of addictive substances in rats

**DOI:** 10.1101/2023.12.14.570833

**Authors:** Ryan D. Farero, Nathan A. Holtz, Suhjung J. Lee, Lauren C. Kruse, Jeremy J. Clark, Paul E. M. Phillips

**Affiliations:** Graduate Program in Neuroscience, University of Washington, Seattle, WA 98195; Center for Neurobiology of Addiction, Pain & Emotion, University of Washington, Seattle, WA 98195; Department of Psychiatry & Behavioral Science, University of Washington, Seattle, WA 98195; Department of Pharmacology, University of Washington, Seattle, WA 98195

**Author notes:** Correspondence to Paul E. M. Phillips.

## Abstract

Dopamine transmission is implicated in aberrant behaviors associated with substance use disorders. Previous research revealed a causal link between excessive drug consumption and the loss of dopamine signaling to stimuli associated with psychostimulant use. The emerging change in dopamine signaling is specific to stimuli associated with the substance rather than the pharmacological properties of the drug itself. Because the change in dopamine signaling was specific to the associated stimuli and not the pharmacological properties of the substance, we examined if treatment with the dopamine precursor, L-DOPA, alters alcohol and opioid self-administration. Therefore, we trained rats to orally self-administer ethanol or the synthetic opioid fentanyl and found that treating animals with L-DOPA significantly reduced consumption of both alcohol and fentanyl. These data suggest dopamine signaling has a vital role in mediating the amount of drug animals will voluntarily take, across multiple classes of drugs. Importantly, these data are preclinical demonstrations of L-DOPA being utilized as a harm reducing treatment in substance use disorders.

## Introduction

Drug addiction is a chronic, relapsing disorder distinguished by compulsive and continued drug seeking, and long-lasting changes to the brain (Koob & Volkow, 2016; NIDA, 2018). Substance use disorders (SUDs) are characterized by the class of drug being abused (APA, 2013), and two classes of substances that have heavily impacted society are alcohol and opioids. A majority of approved pharmacological based treatments for SUDs are specific to the class of drug being abused (e.g., alcohol, opioids, nicotine) (NIDA, 2018). As such, a treatment for SUDs targeting the aberrant physiological changes from repeated conditioning could be beneficial, in treating SUDs of multiple drug classes.

Alcohol and opioids increase dopamine activity in the mesolimbic dopamine system (Di Chiara & Imperato, 1988; Koob & Volkow, 2016), an effect linked to their reinforcing and rewarding properties (Perry et al., 2014). Given the opportunity, animals will administer drugs of abuse to maintain a concentration of extracellular dopamine (Wise, Newton, et al., 1995; Wise, Leone, et al., 1995). Stimuli that are repeatedly paired with drug delivery acquire the ability to increase in extracellular dopamine within the nucleus accumbens (NAc) (Katner & Weiss, 1999; Phillips et al., 2003; Stuber et al., 2005; Day et al., 2007; Flagel et al., 2011). However, it was observed that under conditions of extended drug access, animals who escalate their consumption of cocaine across days show a decrement in phasic dopamine signaling to cocaine-paired cues (Willuhn et al., 2014). In this experiment, phasic dopamine release in the NAc was replenished through systemic L-DOPA/benserazide administration, which also decreased animals’ cocaine intake back to pre-escalated levels (Willuhn et al., 2014). Although pharmacological mechanisms between cocaine, alcohol, and fentanyl differ, we hypothesize that similar allostatic processes are responsible for the decrement of cue-evoked phasic dopamine observed by Willuhn and colleagues (Willuhn et al., 2014), and can be generalized across other drugs of abuse. Therefore, we tested the effects of systemic L-DOPA/benserazide on rodent self-administration of alcohol and fentanyl.

## Methods

### Subjects

A total of 64 Wister rats (18 female, 46 male) from Charles Rivers (Hollister, CA, USA) were used. Animals weighed between 145-165 g (female) and 275-300 g (male) at the beginning of all behavioral experiments. All animals were individually housed in ventilated cage racks and experienced a 12-hour light/dark cycle. All food and water were provided *ad libitum*. Twenty-eight of the 46 male rats underwent alcohol self-administration experiments, however 5 were not used for analysis as they failed to self-administer at least 0.6 g/kg of ethanol (∼approximately 10 operant responses). A total of 12 animals, 6 per sex, underwent two-bottle choice self-administration of fentanyl. An even sex split of 16 animals underwent operant self-administration of fentanyl. The remaining 8 animals, 4 per sex, underwent operant water self-administration.

### Alcohol Self-administration

In addition to the water bottles provided in animals’ home cages, they were given 24-hour access to a second bottle containing an ethanol solution for 20 consecutive days. During the first 7 days of this 20-day period the ethanol bottle contained a 10% ethanol solution, whereas for the final 13 days animals were provided with a bottle containing a 20% ethanol solution. Following the 24-hour two bottle choice paradigm, animals received 3-5 30-minute magazine training sessions before beginning the operant self-administration stage of the experiment. Within this stage, animals were given 25 days of 1-hour access to nose-poke for delivery of 0.2 ml of 20% ethanol. Within this operant task, both an active and inactive nose-poke port were available. A nose-poke in the active port elicited a 5-s presentation of an audiovisual stimulus (nose-poke light + tone) and then a 10-s presentation of the magazine light paired with activation of a solenoid (Lee Valves, West Brook, CT, USA) and consequent liquid ethanol delivery into the magazine bowl. A diagram of the cue-configuration can be found in Figure 1A. Inactive nose-pokes were recorded but not reinforced. Following the 25 days of access, animals then received L-DOPA or saline I.P. injections. If animals failed to reach an average of 0.6 mg/kg of ethanol consumed (∼10 nose-pokes) during the baseline operant sessions (20-25), they were removed from the study. This amount of intake was estimated to be about 0.02% blood alcohol content (Holgate et al., 2017).

**Figure 1.**
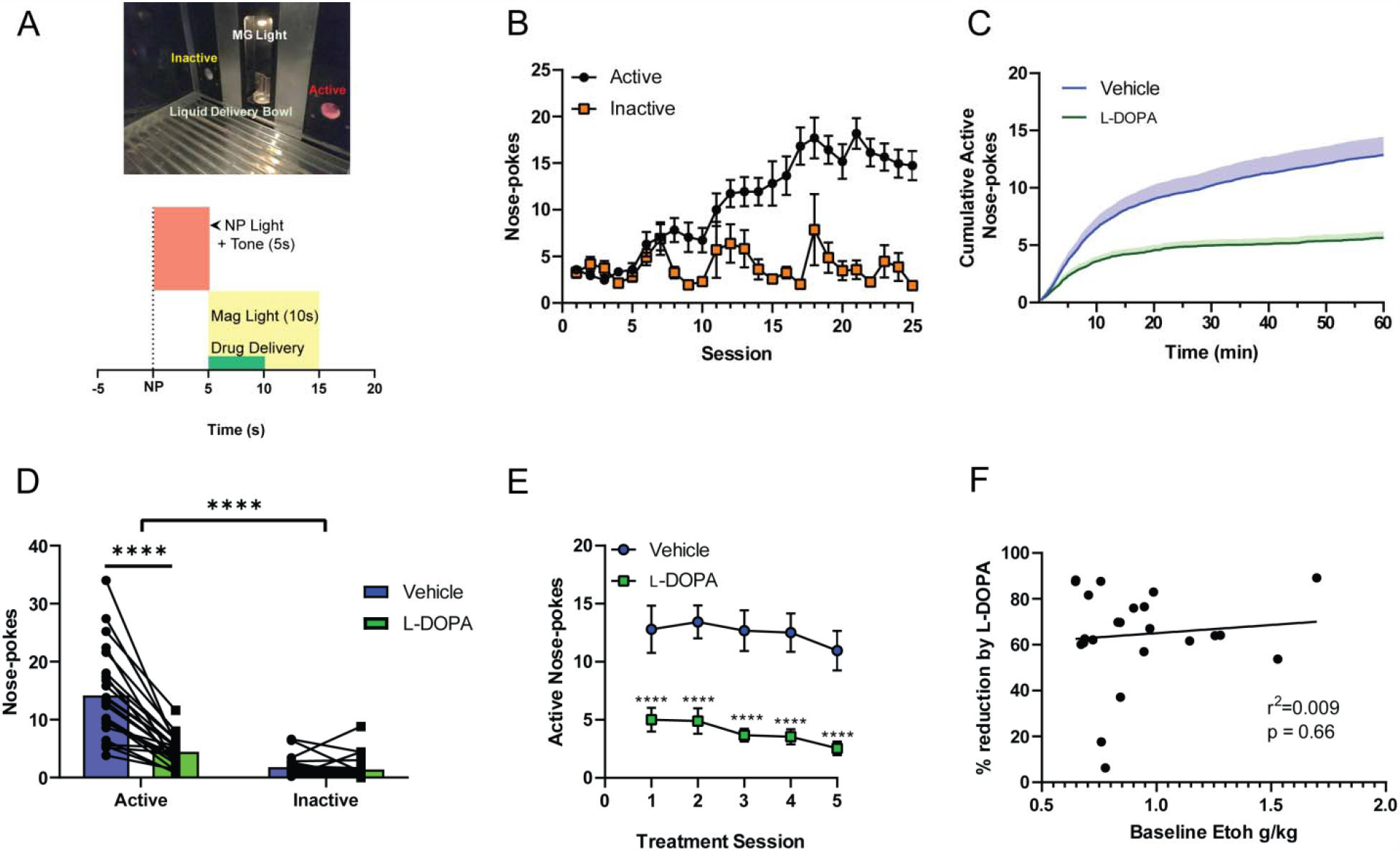
L-DOPA decreases ethanol consumption. **A)** Image of the operant chamber (top panel) and a schematic of the audiovisual stimulus paradigm when an animal performs an active nose-poke. **B)** Animals show a clear discrimination between active and inactive nose-pokes during the 25 sessions provided before any treatment tests. A significant interaction was observed between time and nose-poke port (F(24,528)=8.122, p<0.0001). **C)** Animals were treated with L-DOPA or vehicle, cumulative nose-pokes were calculated and plotted. There is a clear visual difference in active nose-pokes across the session. **D)** The sum of animals active and inactive nose-pokes during treatment sessions were averaged. A two-way ANOVA revealed an interaction of nose-pokes between nose-poke type by treatment (F(1,22)=44.07, p<0.0001). A Sidak post-hoc test, revealed a significant difference in vehicle and L-DOPA active nose-pokes (p<0.0001). **E)** Animals were treated with L-DOPA and vehicle for 5 serial sessions, we analyzed how repeated treatment effects active nose-pokes. A two-way ANOVA found a main effect of treatment (F(1,22)=48.51, p<0.0001). A Sidak post-hoc analysis revealed a significant difference for each treatment day compared to the matched vehicle session (p<0.0001). **F)** A regression of baseline ethanol, calculated from sessions 20-25, and the % of reduction completed by L-DOPA was completed. We found that baseline ethanol intake did not predict the amount of reduction in drug intake from L-DOPA.

### Fentanyl Two-Bottle Choice

In a separate experiment, animals had access to liquid fentanyl (Fagron, Rotterdam, Netherlands) in addition to deionized (DI) water in separate bottles. Fentanyl and DI water were made available to orally self-administer for three hours a day (between 9 a.m. and 5 p.m.). All animals were subjected to a total of 18 sessions before starting treatment/testing sessions. Body weight was recorded before each experimentation session. Fluid consumption of water (DI H2O) and fentanyl solution (50 μg/mL) dissolved in DI H2O was determined by weighing the drinking bottles. The positions of the fentanyl bottles were consistent throughout the entirety of the experiment and counterbalanced between animals in the left and right positions of the cage.

### Oral Fentanyl Operant Self-Administration

Animals learned to self-administer liquid delivery in a modular operant chamber (Med Associates, VT, USA) equipped with two nose-poke devices (port with integrated cue lights) located on adjacent panels of the same wall, a magazine light over a ∼5 mL bowl liquid dispenser stationed between nose ports, a house light, and speakers that emitted a pure-tone. The operant chamber was placed within a sound-attenuated outer chamber. Rats were trained to obtain liquid fentanyl that was delivered into the dispenser following an operant response on FR1 reinforcement schedule. The dispenser was initialized to deliver 0.04 mL/kg of animal’s weight. A nose poke in the active port (side counterbalanced between animals) immediately activated a solenoid valve (Lee Valves, West Brook, CT, USA) to deliver liquid into the dispenser, and was paired with a 10-second presentation of an audiovisual stimulus. The audiovisual stimulus consisted of the illumination of a red light inside the nose-poke port for 1-second, contingent with a the simultaneous presentation of a 10-second tone and magazine light (conditioned stimulus, CS). During CS presentation, a 10-second time out period was imposed during which nose poking did not result in additional drug delivery or any other programmed consequences. Drug availability during the session was signaled by illumination of the house light (discriminative stimulus, DS). Refer to Figure 3 for cue-configuration diagram. To monitor response specificity, nose-poking of the second (inactive) port was recorded but was never reinforced. Rats were given daily access to either liquid fentanyl (50 μg/mL, 0.02 mg/kg/delivery) or DI H2O (0.04 ml/kg/delivery) for one-hour per session over 24-31 days. A subset of animals (4 male, 4 female) were trained to self-administer 0.01 mg/kg/delivery of fentanyl for 13 sessions before increasing the fentanyl dose to 0.02 mg/kg/delivery for another 8 sessions prior to testing sessions. These 8 animals were not included in the analysis or data shown in Figure 3B. The mg/kg and mL/kg was calculated by measuring the amount of liquid left at the end of every session and subtracting from the total amount of liquid delivered.

**Figure 2.**
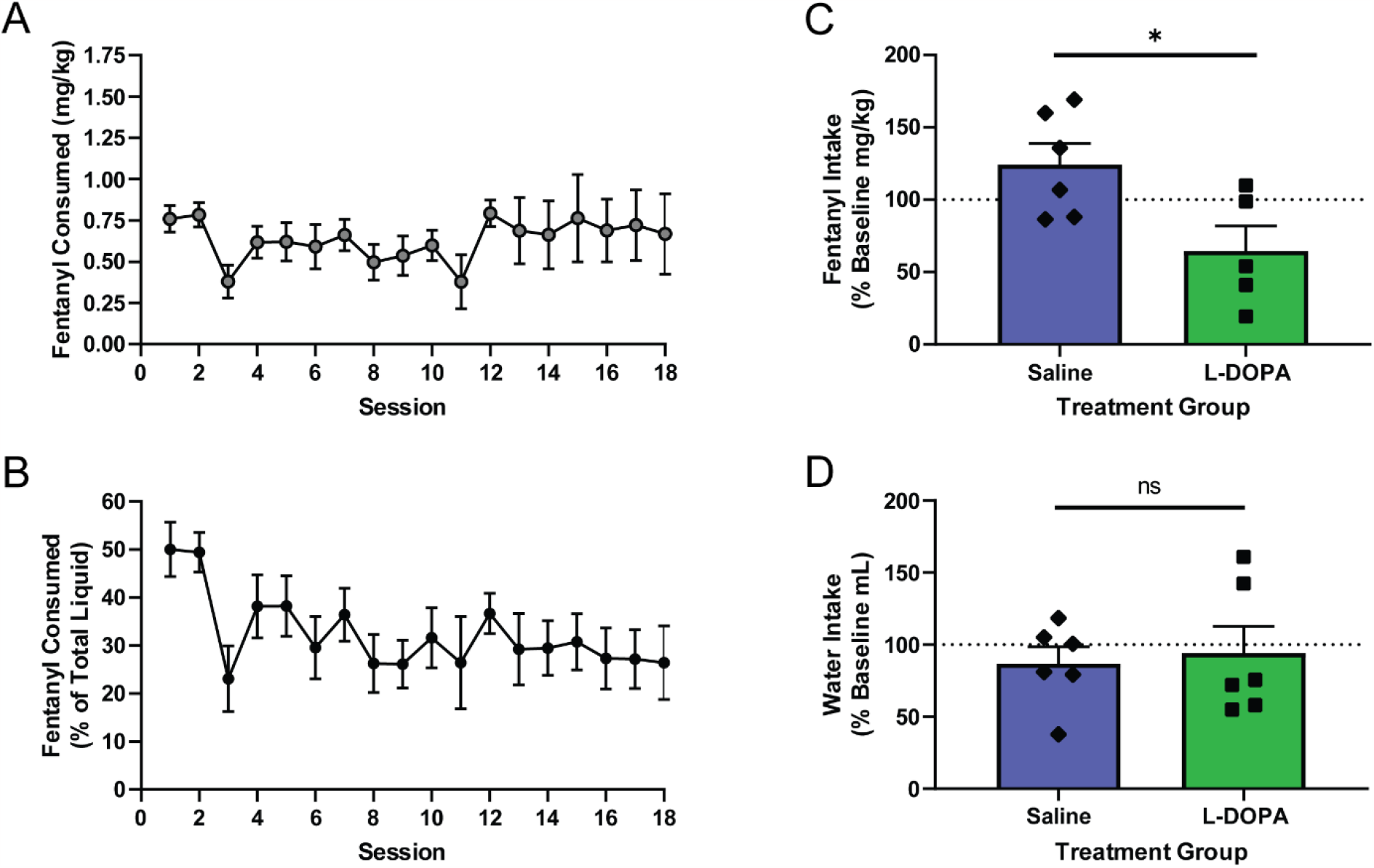
L-DOPA decreases fentanyl but not water consumption in two-bottle choice. **A)** Animals were provided with 18 3-hour sessions to access a fentanyl bottle within their home cage and drank about 0.63 mg/kg of fentanyl. **B)** Within these 3-hour sessions approximately 32% of their total volume intake was from the fentanyl bottle. **C)** Animals were treated with either L-DOPA or saline. Animals treated with L-DOPA had a significant difference in their change of intake from baseline (sessions 15-18) compared to saline treated animals (t(9)=2.658, p<0.05). **D)** The change in amount of water consumed from baseline sessions (15-18), was no different between saline and L-DOPA treated animals (t(9)=0.6189, p>0.05)

**Figure 3.**
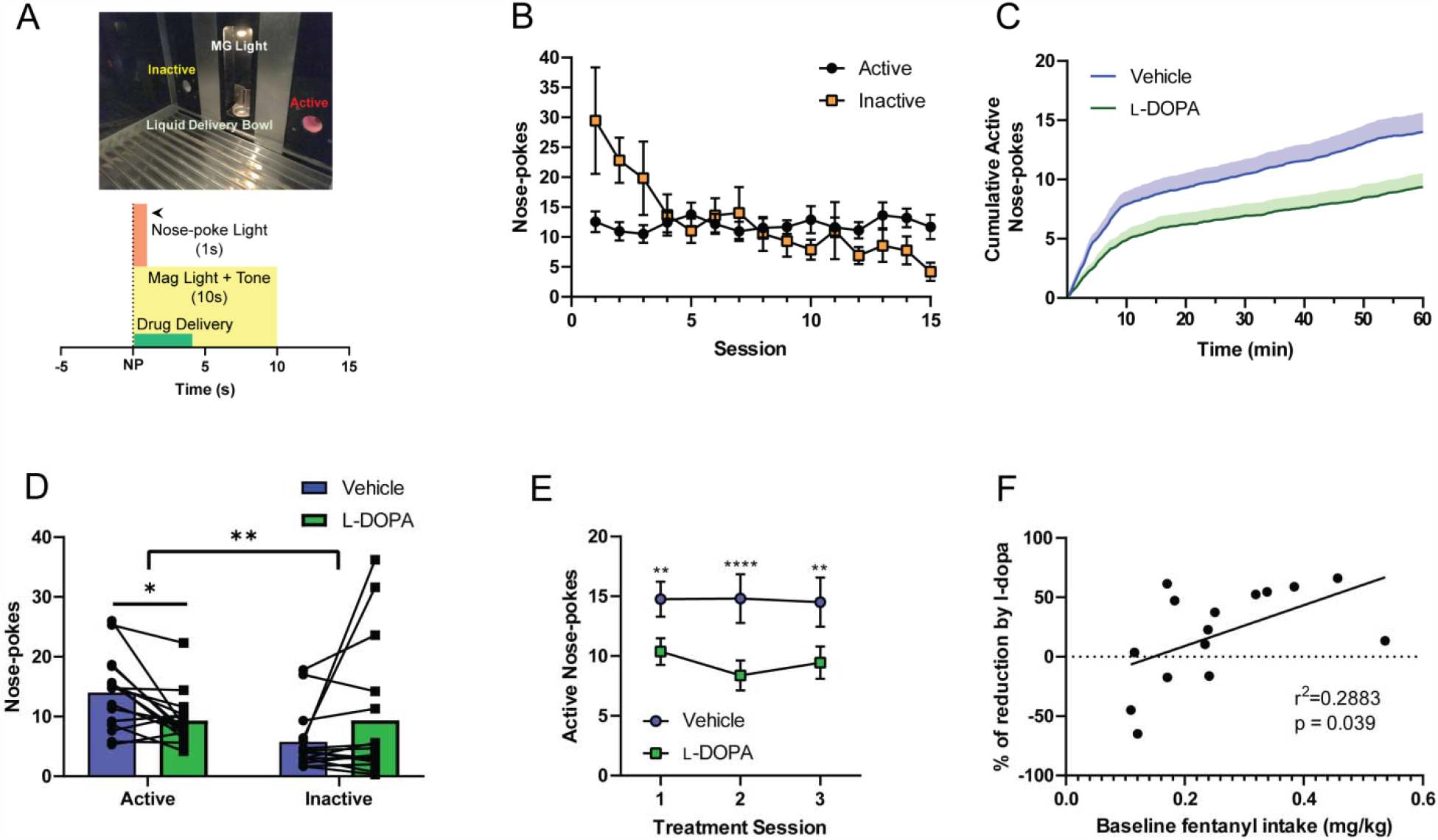
L-DOPA decreases fentanyl responding in an operant self-administration task. **A)** Schematic providing the cue configuration used for oral fentanyl self-administration. **B)** Animals were provided with 15 1-hour sessions with access to fentanyl. A significant interaction was observed between time and nose-poke port (F(14,210)=4.341, p<0.0001). **C)** Animal’s cumulative active nose-pokes were graphed and a clear separation of active nose-pokes between Vehicle treated and L-DOPA treated sessions can be observed. **D)** A comparison of the effect L-DOPA had on active and inactive nose-pokes was performed. Each bar represents the mean with matched data points. A two-way ANOVA revealed a significant interaction between nose-poke type and treatment (F(1,15)=13.32, p<0.01). A Sidak post-hoc test revealed a significant decrease in active nose-pokes during L-DOPA treatment compared to vehicle treatment (p<0.01). **E)** Animal’s received 3-5 days of each treatment (vehicle or L-DOPA). A comparison of individual treatment days was performed. A two-way ANOVA was calculated and found a main-effect of Treatment (F(1,15)=8.877, p<0.01). A Sidak post-hoc analysis was performed to see if L-DOPA treated days were significantly lower than the matched vehicle days. **F)** A linear regression was calculated on baseline fentanyl intake, which was defined as sessions 10-15, and the percent of reduction achieved by L-DOPA. A significant regression was found and highlighted that high baseline fentanyl intake predicted a larger percent reduction in drug-consumption by L-DOPA (r^2^-0.2883, p<0.05). *p<0.05, **p<0.01, ***p<0.001, ****p<0.0001. In B and E, each point represents mean ± SEM.

### L-DOPA/benserazide Administration

Administration of L-DOPA (L-3,4-dihydroxyphenylalanine) (Sigma Aldrich) was paired with the peripherally acting DOPA decarboxylase inhibitor benserazide (Sigma Aldrich) to decrease peripheral breakdown of L-DOPA, and allowing it to reach the brain. Both L-DOPA (30 mg/kg) and benserazide (15 mg/kg) were dissolved in saline and injected intraperitoneally (IP) at a volume of 1 mL/kg body weight 20 minutes prior to experimentation session. All animals in operant self-administration tasks received I.P. injections of both L-DOPA/benserazide and saline. Each treatment was given in 3-5 consecutive days and the order in which saline or L-DOPA was given was counterbalanced between animals. For two bottle choice, L-DOPA/benserazide treatment was injected IP and given before sessions 19-22 of the experiment. Three females and three males received L-DOPA whereas the other three males and three females received saline control.

### Analysis

Simple comparisons between two conditions were performed with unpaired t-tests or paired, depending on the between or within-subject design. Analysis of correlations between variables was conducted using linear regression. In the case of analyses between multiple groups with repeated-measures, ANOVAs were performed using a linear mixed effect modeling approach to account for missing data at random. The model was estimated using restricted Maximum Likelihood (REML). Holm-Sidak correction was used to adjust for post-hoc multiple comparisons. All statistical tests were conducted using GraphPad Prism.

## Results

### L-DOPA Decreased Self-Administration of Ethanol

Animals learned to self-administer 20% ethanol in an operant self-administration task. Once animals interacted with the “active” nose-poke port, an audiovisual stimulus would indicate a successful nose-poke lasting for 5-seconds and immediately followed by the onset of the magazine light and the activation of the solenoid leading to drug delivery (Figure 1A). Prior to this operant task, animals received 20 days of 24-hour access to an ethanol bottle in their home cage (Figure S1), and then up to 5 magazine training sessions. Animals showed a significant increase in their active nose-pokes over sessions, whereas their inactive nose-pokes did not change across the 25 pre-treatment operant sessions (Figure 1B). After receiving 25 sessions with 20% ethanol access, an additional 10 sessions were provided, 5 of which animals were pretreated with a with intraperitoneal injections of L-DOPA and benserazide (30mg/kg and 15mg/kg, respectively). During another 5 sessions animals were treated with saline, and treatment order was counterbalanced. Animals treated with L-DOPA decreased the number of nose-pokes performed resulting in ethanol delivery (Figure 1D). A visual difference in the cumulative active nose-pokes between vehicle and L-DOPA treatment days can be observed within the first 10 minutes of the session (Figure 1C). There was no significant difference in the number of inactive nose-pokes the animals made in treatment vs vehicle days (Figure 1D). This effect persisted across all treatment days (Figure 1E). Previous work (Willuhn et al., 2014) demonstrated that only animals that escalated cocaine consumption had a decrease of drug-intake from L-DOPA administration. We decided to investigate if there was a relationship between baseline ethanol consumption and change in intake after L-DOPA. No correlation (r2=0.009, p=0.66) was observed between how effective L-DOPA was in reducing active nose-pokes and the animals’ ethanol intake during the 5 sessions just prior to the treatment sessions (i.e., sessions 21-25) (Figure 1F).

### L-DOPA Decreased Fentanyl Intake but not Water Intake in a Free Operant 2-Bottle Choice Task

Rats underwent a two-bottle choice task in which they had access to fentanyl (50 μg/ml) and DI water for three hours every day. All animals had food and water *ad libitum* during this 3-hour period as well as the other 21 hours of the day. Animals were given 18 3-hour sessions before receiving any treatment. The average consumption during each session was approximately 0.6572 mg/kg of fentanyl (n=12 rats) (Figure 2A). Because there was no significant sex difference in fentanyl consumption (Figure S2), data from female and male rats were combined for all analyses. During the 18 sessions, the animals’ total liquid consumption within the 3-hour period was ∼32% fentanyl (Figure 2B). A between subject’s design was used to test if L-DOPA/benserazide administration decreases volunteered consumption of fentanyl. One group received I.P. injections of vehicle (n=6, 3 males & 3 females) 20 minutes prior to the 3-hour behavioral session, the other group was administered with 30mg/kg L-DOPA and 15mg/kg benserazide (n=5). Each group received injections for 4 consecutive sessions. The effect of the treatment was compared to the 4 sessions prior to treatment administration (baseline). The group that received L-DOPA treatment consumed significantly less fentanyl from baseline compared to the saline treated group (Figure 2C). Treatment of L-DOPA had no significant effect on water consumption (Figure 2D).

### L-DOPA Decreased Fentanyl Consumption and Operant Responding in a Fentanyl Self-Administration Task

A total of 16 male and 16 female rats received at least 15 sessions of an oral fentanyl self-administration task before being administered L-DOPA/benserazide, in order to test its effect on self-administration. Once animals interacted with the active nose-poke port, the LED lights within the nose-poke port would turn on for 1s to indicate a successful nose-poke. Simultaneously, an audiovisual stimulus (magazine light and a tone) would turn on for 10s, and this was also paired with drug delivery through the opening of a solenoid valve (Figure 3A). While animals learned this task, they showed no statistically significant increase in nose-pokes over the course of 15 sessions. However, there was a decrease in the number of inactive nose-pokes being performed, and we found a significant interaction between sessions and nose-poke port (Figure 3B). These animals received no magazine training or prior experience to a 2-bottle choice paradigm such as those in the ethanol self-administration task (Figure 1). In sessions 16-25, animals were administered both L-DOPA/benserazide (30mg/kg:15mg/kg) and saline IP injections for 3-5 sessions, with order being counterbalanced between animals. During sessions in which L-DOPA was administered, there was a visible change in the slope in the cumulative active nose-poke record at the beginning of the session (Figure 3C). A significant interaction between nose-poke type (i.e. active and inactive) and treatment was observed. Sidak p*ost-hoc* analyses revealed that there was a significant decrease in active nose-pokes, whereas no significant change was found in nose-pokes at the inactive port (Figure 3D). Looking at all animals’ first 3 treatment days (half the animals received 5 treatment days, data not shown), there was a main effect of treatment (L-DOPA vs saline). Sidak post-hoc analyses revealed that each day was significantly different from their matched vehicle treated sessions (Figure 3E). We examined if animals’ baseline intake (consumption during sessions 10-15) correlated with L-DOPA-induced decreases in fentanyl consumption. There was a significant regression observed (r^2^=0.2883, p=0.039), in which animals with lower intake of fentanyl during sessions 10-15 had less reduction in intake during L-DOPA treated days (Figure 3F). Animals did not drink all the liquid in the bowl from every session, and at the end of each session any remaining liquid was accounted for and subtracted from the amount delivered. Performing this allowed us to calculate the amount of fentanyl consumed. In Figure S3, this data is presented and shows that not only did animals treated with L-DOPA respond less for fentanyl delivery, but they also drank less fentanyl as well.

### L-DOPA Decreased Operant Responding for Water

Seeing that systemic L-DOPA administration was therapeutically effective in decreasing fentanyl and alcohol consumption, we tested if this effect was specific to drugs of abuse by having animals self-administer water. 4 male and 4 female rats underwent a similar paradigm as that of fentanyl operant self-administration where instead water was delivered at 0.4 mL/kg per active nose poke. Animals readily nose-poked preferentially at the active port (Figure 4B) for delivery of water. Treating animals with L-DOPA had a similar effect on water self-administration compared to fentanyl and alcohol self-administration (Figures 4C, D, & E). L-DOPA/benserazide and saline IP injections were given to each animal 5 sessions for each treatment. Similarly, to ethanol and fentanyl, t visualWe found a separation of cumulative active nose pokes between saline and L-DOPA treatment as shown in Figure 4C, which suggested that. L-DOPA also significantly decreased both average active nose pokes but had no significant effect on inactive nose-pokes (Figure 4D). There was no correlation (r2=0.341, p=0.128) between animals’ baseline operant self-administration of water and the effect L-DOPA had on water consumption (Figure 4F).Tdrank is notablethat they*ad libitum* access outside of experimental sessions

**Figure 4.**
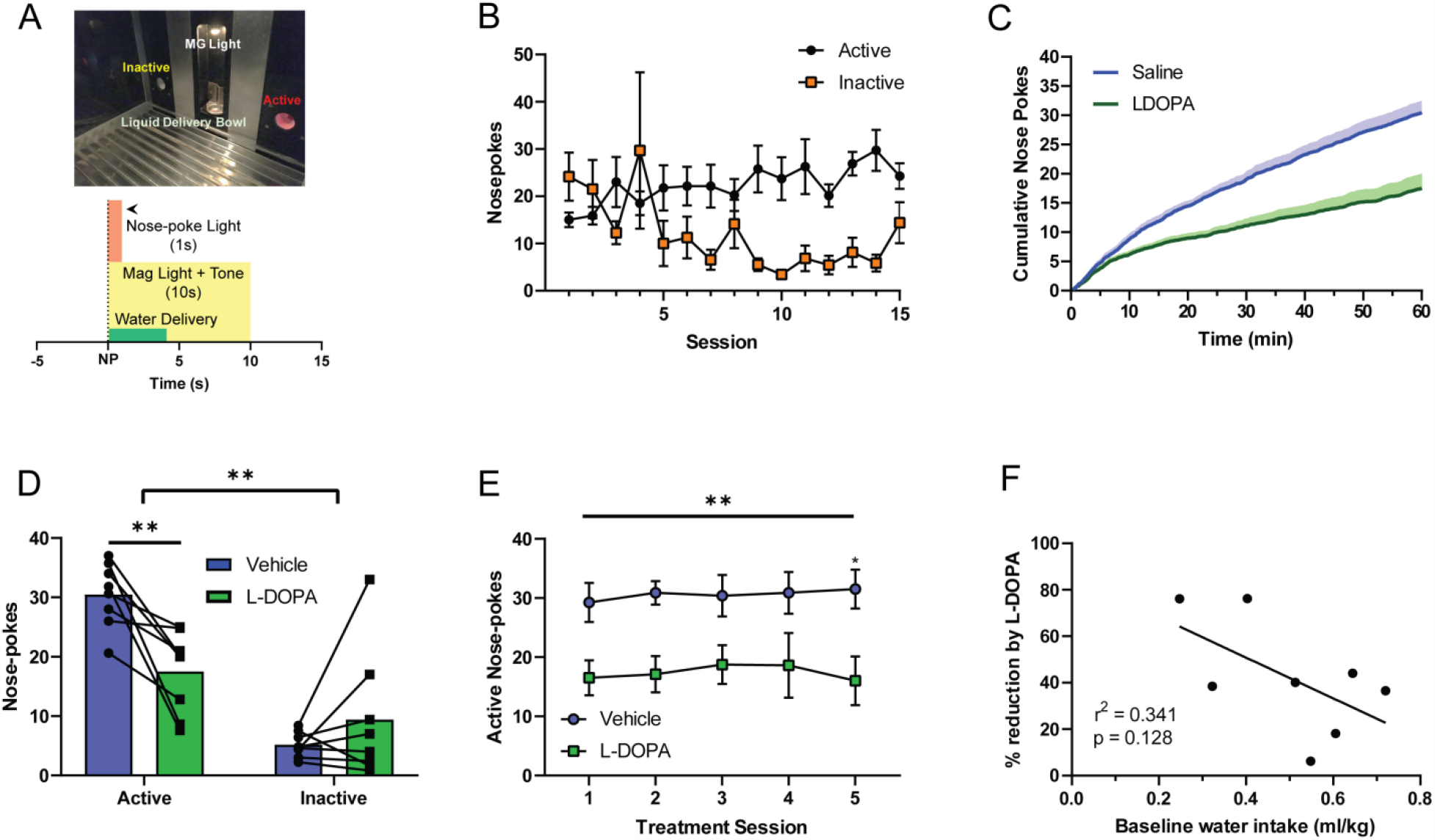
L-DOPA decrease water consumption in an operant task. **A)** Schematic with the CS configuration used during operant water self-administration task. **B)** Animals were given 15 1-hour sessions to self-administer water. An interaction of nose-pokes was observed between sessions and nose-poke type (F(14,180)=4.891, p<0.0001). **C)** Cumulative active nose-pokes were calculated for saline treated and L-DOPA treated sessions. **D)** Total nose-pokes within the 1 hour were used to analyze how L-DOPA affected nose-poking behavior. A two-way ANOVA was performed and a interaction of nose-poke type by treatment was observed (F(1,7)=14.76, p<0.01). Each bar represents the mean with matched data points. A Sidak post-hoc test was completed and revealed a significant difference in active nose-pokes between the two treatment (p<0.01). **E)** The 10 sessions of treatment were separated into matching sessions. A two-way ANOVA revealed a main effect of drug treatment (F(1,7)=16.05), p<0.01). **F)** A linear regression was performed between baseline water intake (sessions 10-15), and % reduction in responding achieved from L-DOPA. Baseline water intake does not have a significant relationship with the reduction of responding by L-DOPA. In B and E, each point represents mean ± SEM.

## Discussion

Our results indicate that L-DOPA administration, given with a peripheral acting DOPA decarboxylase inhibitor, decreased consumption of both alcohol and fentanyl. The underlying model of these experiments hypothesize that diminished cue-evoked dopamine release within the NAc promotes drug consumption, which has been previously shown during psychostimulant self-administration (Willuhn et al., 2014). Given that this treatment has reduced drug-consumption in all classes of drugs we tested, we reason that L-DOPA may be an effective therapy to manage substance use disorders in conjunction with behavioral and/or cognitive therapies.

We established a behavioral paradigm that achieves reliable self-administration of 20% ethanol in male Wistar Rats. Within this paradigm, rodents intake was comparable to that of the “drinking in the dark” paradigm (Holgate et al., 2017). Out of 28 rats, only 5 animals did not reach the criteria of 0.6 g/kg of ethanol consumed during the 1-hour self-administration sessions. Although our results indicate a similar neurochemical change of dopamine transmission’s role in promoting ethanol consumption, we are unaware if the NAc core is the site of action mediating the decrease of drug-consumption. Interestingly, in our ethanol self-administration paradigm, we observed a “load-up” phase (Figure 1C). This is marked by a higher rate followed by a lower stable rate of nose-pokes, similar to that observed in psychostimulant self-administration paradigms (Ahmed & Koob, 2005).

L-DOPA was effective at decreasing fentanyl consumption in two self-administration paradigms. We provided male and female Wistar Rats 3-hour fentanyl access in a two-bottle choice paradigm. We found that animals drank approximately 3 times more fentanyl in the two-bottle choice paradigm than the operant task, which suggests that animals are taking an equal amount of drug per hour, as the two-bottle choice task was also 3 times longer. Within the two-bottle choice task, animals did not show a preference for fentanyl, as only ∼30% of their total liquid intake was from the fentanyl bottle. One interpretation of this outcome is that the animals found fentanyl aversive; however, their intake, after approximately 3 days of drug access, remained stable. Another interpretation of the animals not showing a preference for fentanyl is they were modulating their consumption to maintain a relatively stable drug concentration or hedonic set-point (Wise, Newton, et al., 1995; Wise, Leone, et al., 1995; Ahmed & Koob, 2005). This line of thinking is supported by the fentanyl self-administration operant task, where animals showed a “load-up” phase of active nose-pokes (Figure 3C),

An inactive nose-poke port was utilized as a control operandum in the operant self-administration tasks, and a water bottle within the two-bottle choice task. In all operant tasks, L-DOPA treatment did not significantly alter inactive nose-poke responding. Within the two-bottle choice task, we found that animals treated with L-DOPA did not alter how much water they drank during the 3-hour test period, indicating that L-DOPA reduction is drug specific. However, we ran a control operant experiment in which we trained animals to nose-poke for ∼0.2 mLs of water. Interestingly, when treated with L-DOPA, animals decreased their responding and consumption of water in the operant task, contradicting the results we saw within the two-bottle choice experiment. It is probable that the cue paradigm utilized in the operant task is contributing to the findings of decreased water consumption from L-DOPA, and that these findings would generalize to multiple appetitive cued behavior. Importantly, we only observed the decrease in water intake in an isolated and separate environment from where their normal water and food intake occur. Within their home-cage where the two-bottle choice experiment took place we did not see any effect of L-DOPA on water intake.

One interesting aspect observed by Willuhn and colleagues (Willuhn et al., 2014) wherein L-DOPA was administered to animals during cocaine self-administration, a decrease is drug consumption was observed only with animals that had escalated their intake over sessions, whereas L-DOPA had no effect on animals that did not escalate their intake. Thus, they saw a bimodal distribution of animals that were affected by treatment and those that were not. They classified these animals based on their drug-taking patterns and provided them with prolonged drug access (Willuhn et al., 2014), which produces escalated intake over time (Ahmed & Koob, 2005). To assess if L-DOPA’s effects were related to previous drug or water consumption levels, we used the average intake of each individual animal the week before treatment occurred and compared it to the percent decrease in consumption from vehicle to L-DOPA treated sessions. Surprisingly, only animals in the operant fentanyl task showed a significant relationship between baseline intake and effectiveness of L-DOPA. One explanation for only finding this in the fentanyl group is that animals had only experienced fentanyl for 15-21 days prior to testing days, whereas in alcohol studies they were introduced to ethanol for 20 days in a two-bottle choice paradigm and then another 25 operant sessions. Given the different histories of experience, it is likely that the average intake the 5 sessions before testing is not an accurate baseline.

In conclusion, these experiments provide foundational preclinical evidence that L-DOPA treatment has potential to be used as a harm-reduction therapy for SUDs. In a clinical trial, L-DOPA was paired with contingency management therapy in cocaine abusers, and it was found to prolong abstinence for longer periods compared with placebo (Mariani & Levin, 2012). Although here we showed that L-DOPA decreases intake, rather than prevents intake, L-DOPA may have multiple uses in treating persons living with SUDs.

## Acknowledgements

This work was funded by the National Institutes of Health grants F31-DA048562 (RF), T32-AA007455 (PP), R01-AA021121 (JC), U01-AA024599 (JC), R01-DA039687 (PP), and R37-DA051686 (PP).

## Supplemental Figures

**Figure S1.**
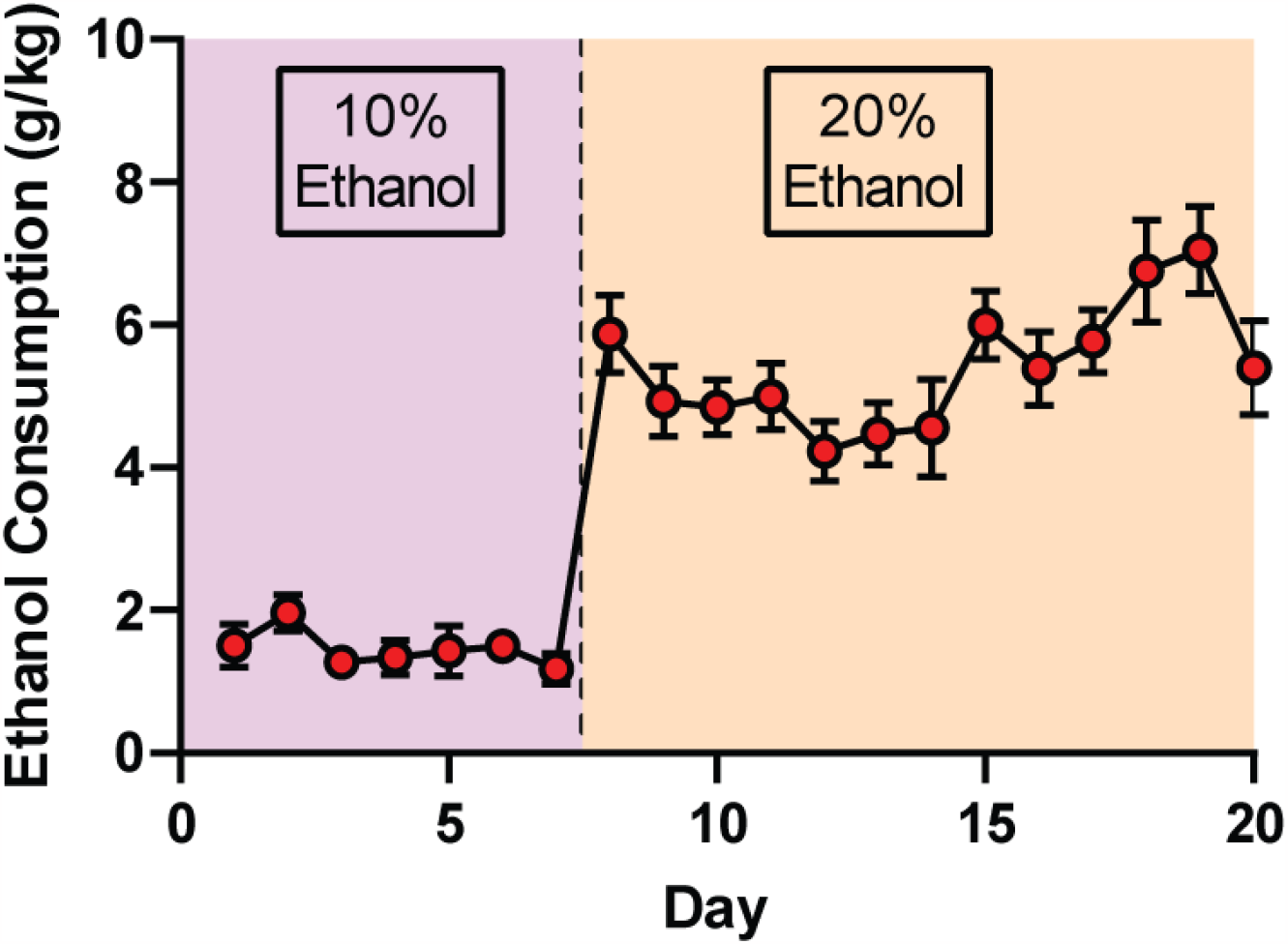
Ethanol consumption during 24-hour two-bottle choice prior to operant training. During the first 7 days animals were provided access to 10% ethanol (pink box). From day 8 to day 20 the concentration of ethanol was increased to 20%. Each point represents mean ± SEM.

**Figure S2.**
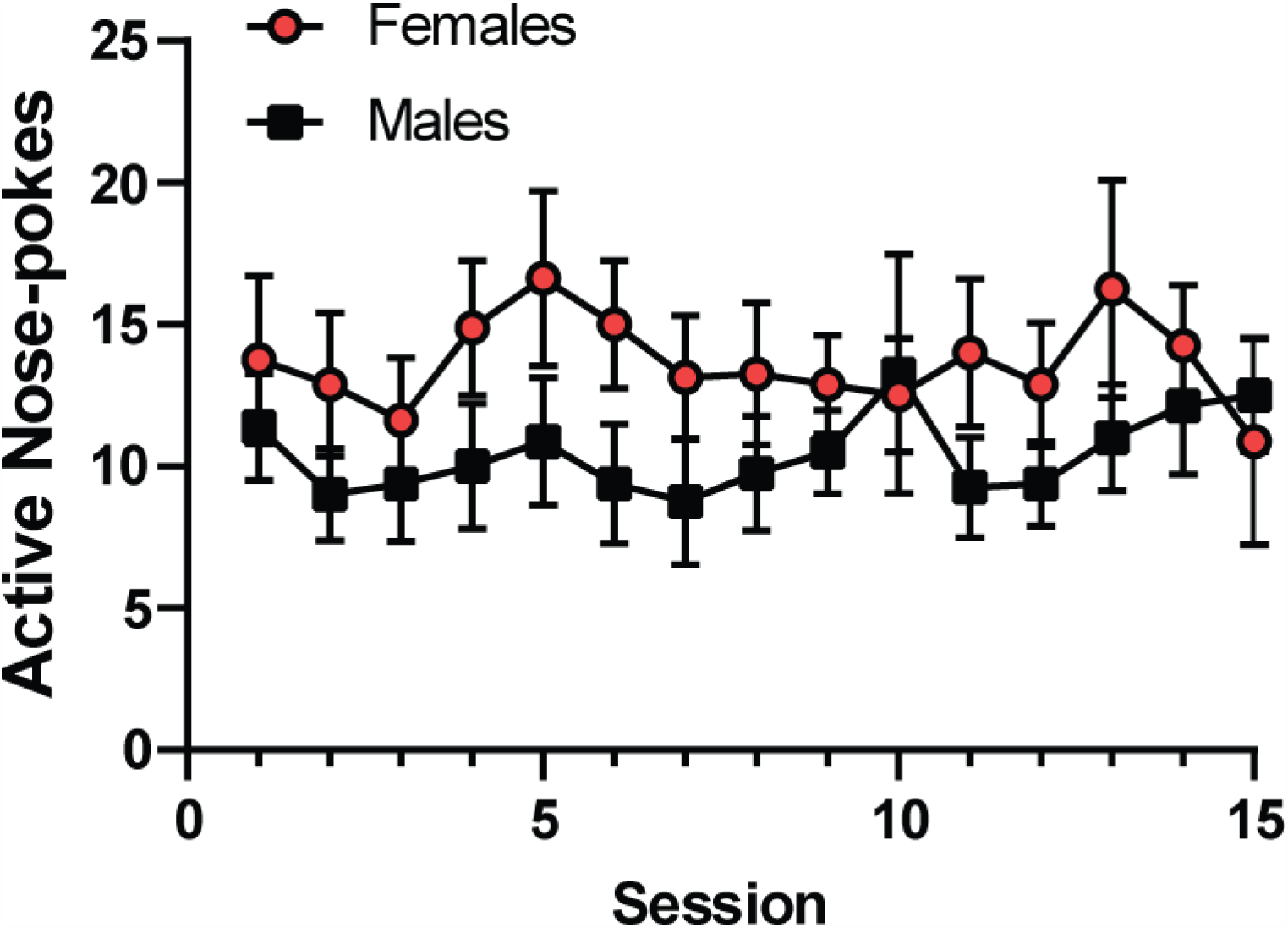
Males and females did not differ in their active nose-poke during the first 15 sessions provided fentanyl self-administration. A two-way ANOVA was completed and found no significant differences in active nose-pokes between sex (F(1,14)=1.573, p>0.05). Each point represents mean ± SEM.

**Figure S3.**
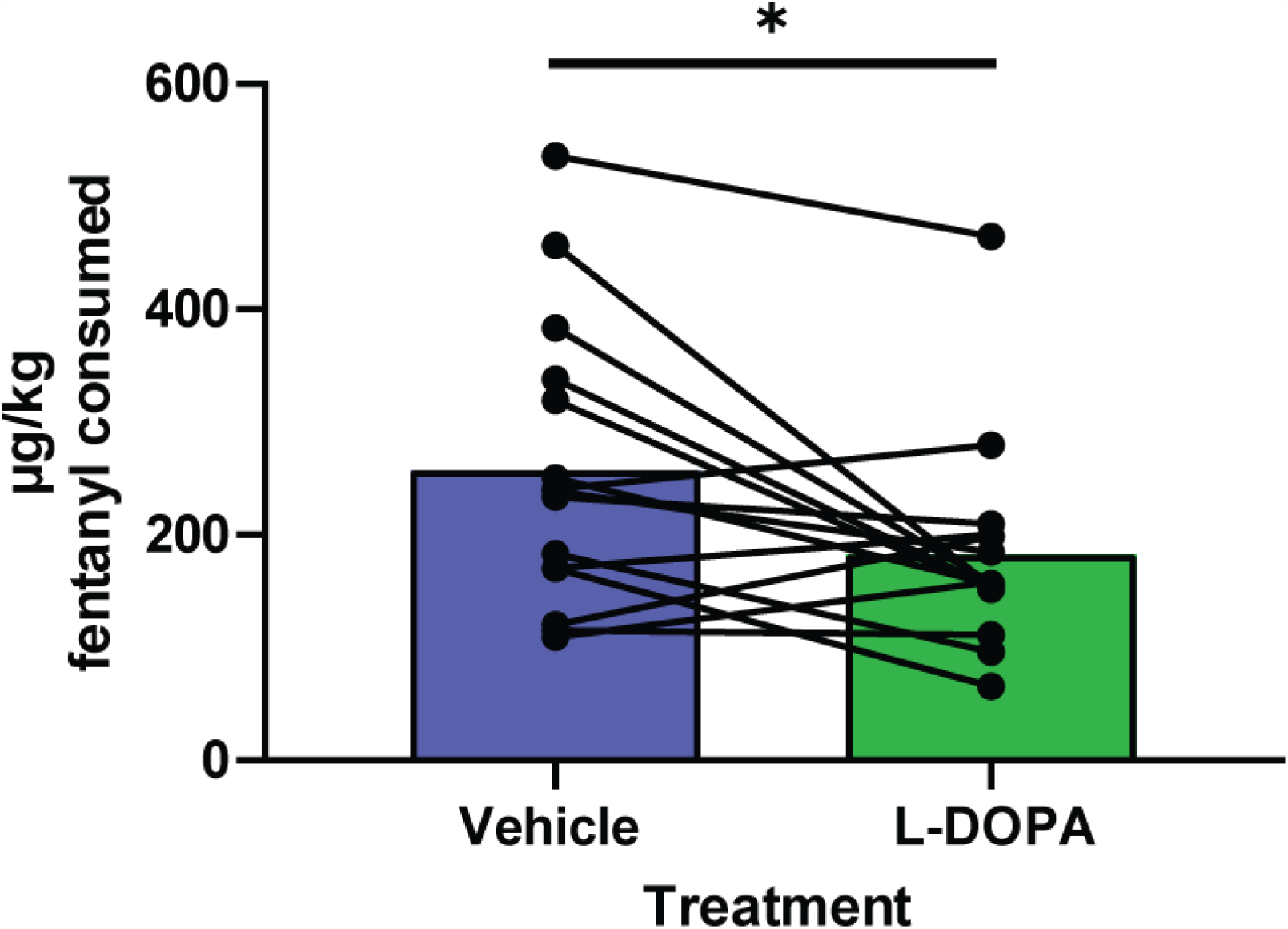
L-DOPA decreased total amount of fentanyl consumed. Each bar represents the mean with matched data points. The amount of fentanyl consumed was calculated by subtracting the amount of liquid left over after the session. A significant decrease of fentanyl consumed was observed (paired t-test, t(14)=2.655, p<0.05).

